# The m6A reader YTHDC2 is essential for escape from KSHV SOX-induced RNA decay

**DOI:** 10.1101/2021.09.03.458900

**Authors:** Daniel Macveigh-Fierro, Angelina Cicerchia, Ashley Cadorette, Vasudha Sharma, Mandy Muller

**Affiliations:** Department of Microbiology, University of Massachusetts, Amherst, USA; Molecular and Cellular Biology Graduate Program, University of Massachusetts, Amherst, United States

## Abstract

The role m^6^A modifications have increasingly been associated with diverse set of roles in modulating viruses and influencing the outcomes of viral infection. Here we report that the landscape of m^6^A deposition is drastically shifted during KSHV (Kaposi Sarcoma Associated herpesvirus) lytic infection for both viral and host transcripts. In line with previous reports, we also saw an overall decrease in host methylation in favor of viral mRNA along with 5’ hypomethylation and 3’ hypermethylation. During KSHV lytic infection, a major shift in overall mRNA abundance is driven by the viral endoribonuclease SOX, which induces the decay of greater than 70% of transcripts. Here, we reveal that Interlukin-6 (IL-6) mRNA, a well-characterized SOX-resistant transcript, is m^6^A modified during lytic infection. Furthermore, we show that this modification falls within the IL-6 SOX Resistance Element (SRE), an RNA element in IL-6 3’ UTR that was previously shown to be sufficient for protection from SOX cleavage. We show that the presence of this m^6^A modification is essential to confer SOX resistance to the IL-6 mRNA. We next show that this modification recruits the m^6^A reader YTHDC2 and found that YTHDC2 is necessary for the escape of the IL-6 transcript. These results shed light on how the host cell has evolved to use RNA modifications to circumvent viral manipulation of RNA fate during KSHV infection.

## Introduction

DNA and RNA viruses regulate the transcriptional and post-transcriptional fate of host mRNA to gain access to key resources during infection. Many diverse viruses, including the gammaherpesvirus Kaposi’s sarcoma-associated herpesvirus (KSHV), trigger a widespread mRNA decay event known as “host shutoff” that decimates greater than 70% of the cellular transcriptome [1-5]. To achieve this level of degradation, KSHV encodes ORF37 (SOX), an endoribonuclease conserved throughout the gammaherpesvirus family. SOX is responsible for host shutoff, using RNA degradation to dampen cellular gene expression and any mounting immune responses [1, 5, 6], allowing the virus access to newly freed host resources for viral replication. SOX is known to internally cleave cytoplasmic mRNAs in a site-specific manner which then promotes degradation by the cellular 5’ to 3’ exonuclease Xrn1 [7-9]. However, over the past decade, we and others have found select mRNA transcripts that robustly escape SOX-induced decay [10, 11]. Multiple mechanisms have been hypothesized to explain what contributes to promoting escape from SOX, ranging from a lack of a SOX targeting motif to indirect transcriptional effects [12, 13]. More recently, we have demonstrated that there is a smaller subset of cellular transcripts that actively evade SOX. These “dominant” escapees each carry a specific RNA element found within their 3’ UTRs termed the <u>S</u>OX <u>r</u>esistance <u>e</u>lement (SRE). The SRE confers protection to the target transcript from SOX even if the transcript contains a SOX targeting motif [10, 14, 15]. Interestingly, the SRE can resist multiple viral endonucleases but not cellular endonucleases, making it a virus-specific RNase escape element [15, 16].

To date, it is still unknown how many of these SREs are present in the genome, their mechanism of action against viral endonucleases, or what becomes of the SRE containing transcripts once they are spared from degradation. So far, three SRE-containing “escapees” have been identified: interleukin-6 (IL-6), growth arrest DNA damage-inducible 45 beta (GADD45B) and C19ORF66 [10, 14-17]. Although there is little sequence homology among known SREs, they share similarities in their secondary structures, bolstering the idea that the SRE may act as a platform for the recruitment of a protective protein complex [15, 16]. Furthermore, it was observed that the SRE is only active when located in the 3’ UTR region of a transcript, suggesting that this RNA element likely functions in conjunction with proteins to modulate RNA stability [14, 15]. Previous mass spectrometry screens have identified several host proteins that can bind to the SRE and intriguingly, a few of these proteins are m^6^A readers.

N^6^-methyladenosine (m^6^A) is the most prevalent mRNA modification out of the over hundred different known RNA modifications [18, 19]. m^6^A impacts virtually every stage of post-transcriptional mRNA fate from splicing, localization, translation, and decay [20-24]. Deposition of m^6^A occurs co- or post-transcriptionally *via* an m^6^A writer complex that consists of a catalytic methyltransferase subunit, such as METTL3, and other cofactors such as METTL14/16 and WTAP [25-29]. The writer complex recognizes a DRACH (D=G/A/U, R=G/A, H=A/U/C) motif for methylation. Although a transcript can have several DRACH motifs, not all will be methylated [25]. What determines which motif will be chosen is still unknown [30, 31]. Demethylases or erasers like FTO and ALKBH5 add a layer of reversibility to the m^6^A epitranscriptome [32, 33]. The presence or absence of these modifications can change the secondary structure of mRNA and create platforms for m^6^A reader proteins. Reader proteins then recognize the modification and promote specific RNA fates in turn [21, 25-29, 34].

Recent transcriptome-wide m^6^A mapping of multiple viruses [35-40] have brought to the forefront research concerning a complex interplay between the m^6^A pathways and viral replication success. Previous research has shown that KSHV can hijack this system to deposit m^6^A onto its own transcripts, including KSHV ORF50 (RTA), the master latent-to-lytic switch protein and on the multifunction long-noncoding RNA (lncRNA) PAN [36, 41-43]. While there is strong evidence that the m^6^A landscape is reshaped during KSHV infection, and that this shifts ultimately promotes the progression of KSHV infection, it remains unclear whether this m^6^A repurposing also affects the fate of host transcripts.

Here we show that the IL-6 mRNA is m^6^A-modified in its 3’UTR during KSHV lytic infection and that removal of this m^6^A mark restores susceptibility to SOX-mediated degradation. We further show that the m^6^A reader YTHDC2 binds to the IL-6 SRE in an m^6^A-dependent manner and that downregulation of YTHDC2 is sufficient to abrogate resistance to SOX. Taken together these results demonstrate that the m^6^A pathway is pivotal in the regulation of gene expression during KSHV infection, highlighting the viral-host battle for control of RNA stability.

## Results

### KSHV infection reshapes the m6A landscape in cells

Since KSHV reactivation broadly affects RNA fate and extensively remodels the host gene expression environment, we hypothesized that m^6^A modifications may be broadly redistributed upon KSHV lytic reactivation from latency. We mapped transcriptome-wide m^6^A modification sites with single nucleotide resolution using m^6^A-eCLIP. RNA was isolated from KSHV positive iSLK.219 cells either in their latent state (Lat) or lytic state 48h post reactivation (Lyt) and an anti-m^6^A antibody was used to enrich m^6^A-modified RNA fragments prior to RNA sequencing of both the input and immunoprecipitated (IP) samples (FIG 1A). We detected a total of 2281 peaks in the latent samples and 1482 peaks in the reactivated samples, corresponding to over 40,000 unique called sites which 54% of sites were identical in both of our samples. As expected, the m^6^A motif DRACH (in particular, [GGACU]) was enriched under the identified peaks, confirming that our m^6^A deposition in these infected cells is concordant with previous observations (Fig 1B) [35, 36]. Also, in agreement with previous data, m^6^A peaks were most prevalent around the transcript STOP codon and beginning of the 3’UTR (FIG 1C). The overall m^6^A peak deposition profiles between Lat and Lyt samples were surprisingly close, however, we observed decreased methylations in cellular mRNA 5’ UTRs upon KSHV lytic reactivation, which is in contrast with observations in other viruses such as ZIKV. GO-term analysis of genes with lytic-specific peaks identified an enrichment for genes with roles in RNA splicing, while genes involved in DNA replication seem to carry less m^6^A modifications in KSHV lytic cells (FIG 1D). We also detected several m^6^A peaks within viral genes, many of which have been characterized before (FIG 1E) [35, 36, 43]. Together, these results reinforce past observations that KSHV infection affects the broader m^6^A profile of cells, redistributing m^6^A modifications to different host and viral genes, which likely has far-reaching consequences on modulation of gene expression.

**Figure 1:**
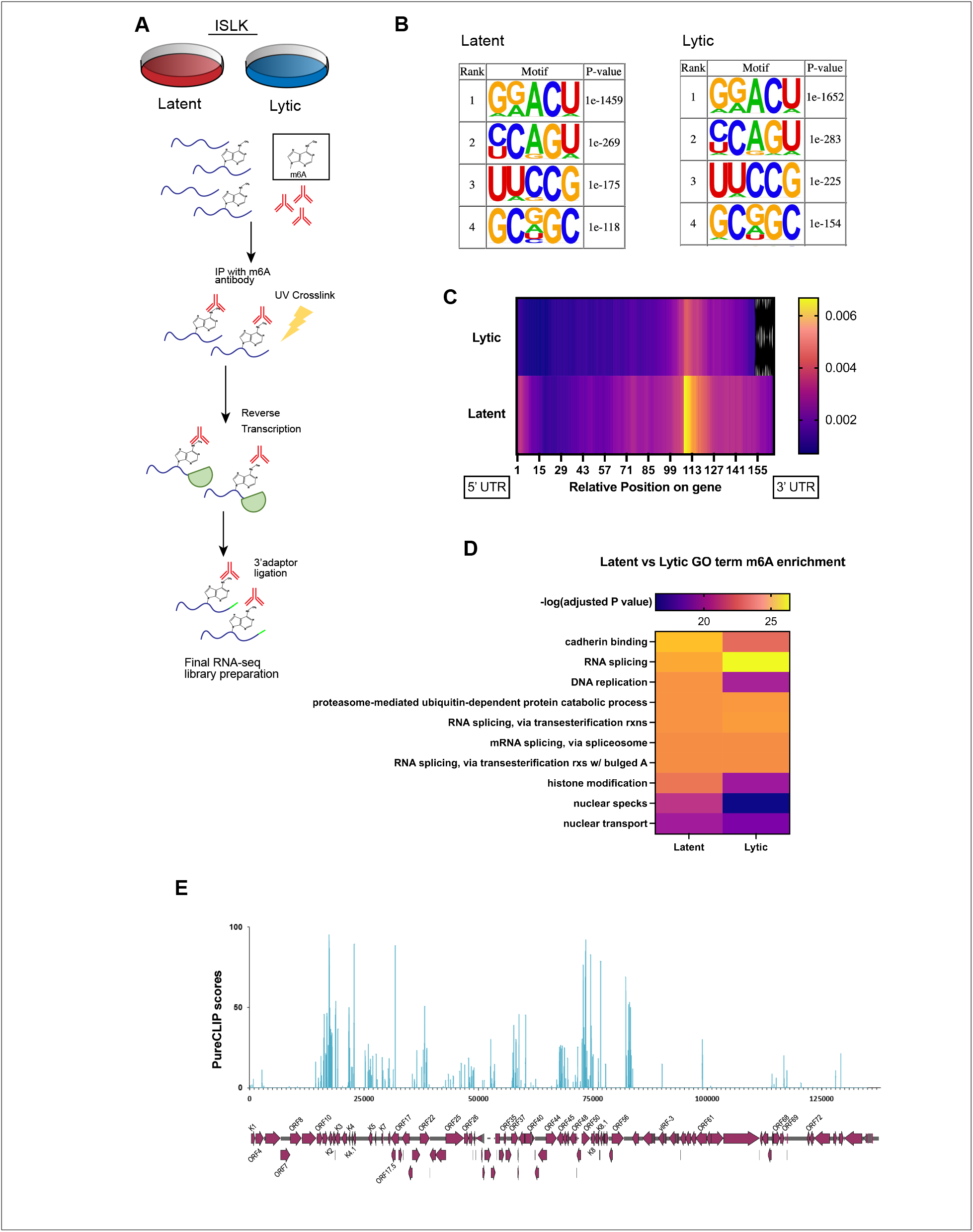
Examining ISLK m^6^A epitrascriptome during KSHV lytic reactivation. **A)** Schematic of m^6^A eCLIP set up. iSLK.WT cells were either left latent or lytically reactivated with doxycycline and sodium butyrate for 48 hours. Total RNA was collected then incubated with an m^6^A antibody. Samples were UV crosslinked before being reverse transcribed then attached with 3’ adapters in part of library preparation. Finally, m^6^A enriched samples were sequenced. **B)** Most significant DRACH motifs with m^6^A peaks identified by HOMER in latent and lytic cells. **C)** Heat map of a metagene plot depicting the average number of sites mapped to certain genomic regions. The number of sites is calculated for each region of every gene, the lengths of the regions are then normalized, and the average number of sites for a set number of positions along the regions are calculated. **D)** Heat map of the most significant m^6^A enriched functional pathways in latent and lytic cells calculated through an enrichment analysis preformed using the R package clusterProfiler. **E)** m6A PureCLIP scores of lytically reactivated KSHV genes aligned over an annotated KSHV genome. PureCLIP is the log posterior probability ratio of the m^6^A crosslink sites over the input samples.

### IL-6 SRE carries a lytic-specific m^6^A modification

We next focused our attention on the interleukin-6 transcript, the best characterized SOX-resistant mRNA. In latent cells, we detected several m^6^A peaks between positions 22,727,199 - 203 which corresponds to IL-6 5’ UTR. However, in lytic cells, IL-6 gains an additional peak at position 22,731,646, corresponding to IL-6 3’ UTR. This m^6^A modification falls on the SRE region and on a strong DRACH motif (FIG 2A). To confirm the presence of this m^6^A deposition, we mutated the predicted position within the SRE (referred to as SRE*) and performed meRIP-qPCR to assess m^6^A deposition on the WT SRE compared to the SRE*. meRIP-qPCR confirmed the presence of the m^6^A peak within the SRE and that mutating nucleotide 74 within the IL6 SRE is enough to abrogate m^6^A pulldown (FIG 2 B).

**Figure 2:**
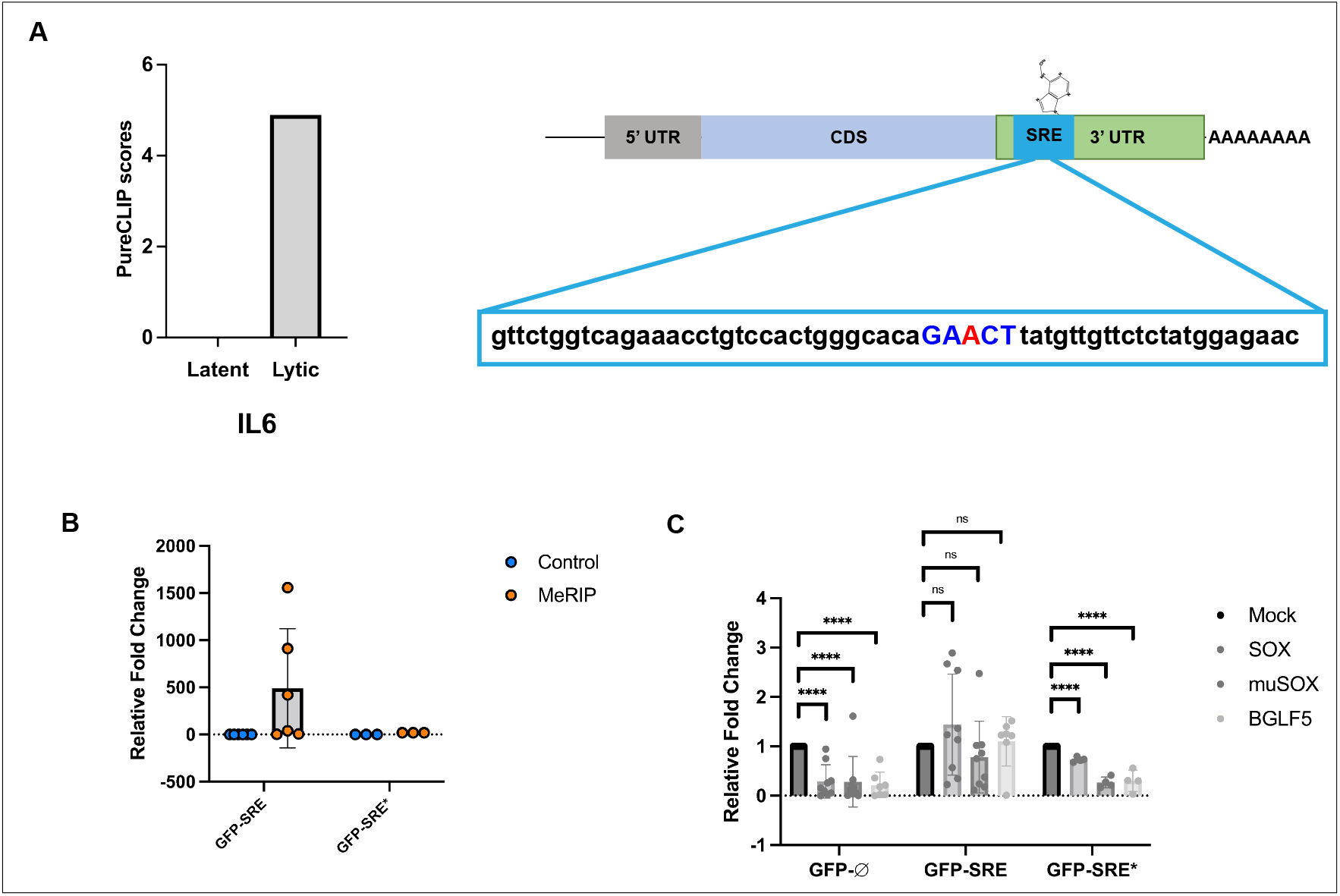
IL-6 SRE contains an m^6^A site that is necessary for viral endonuclease protection. **A)** PureCLIP scores of the 3’ UTR IL-6 gene in latent and lytic (48hpr) ISLK.WT cells. Schematic to the right illustrates an IL-6 gene: the DRACH motif identified through m6A-eCLIP is in blue, and the methylated adenosine in red. **B)** Cells were transfected with WT SRE or SRE* GFP reporter, and total RNA was harvested 24hr later and subjected to meRIP followed by RT-qPCR using GFP primer. Fold enrichment was determined by calculating the fold change of the IP to control Ct values that were normalized through the input. **C)** 293T cells transfected with one of three viral endonucleases as indicated along with the indicated GFP reporters. RNA was collected and quantified using RT-qPCR.

We next investigated whether this m^6^A modification plays a role in SRE-mediated escape from SOX-induced decay. Cells were transfected with a SOX construct (or mock) alongside a GFP expressing reporter bearing no SRE (GFP-Ø) and thus susceptible to SOX, or a GFP reporter fused to a WT SRE (GFP-SRE), expected to be protected from SOX, or fused to SRE* (GFP-SRE*) to test the effect of the loss of m^6^A deposition on escape from SOX. As shown in Fig 2C, as expected, SOX efficiently degrades GFP-Ø but GFP-SRE resists degradation. However, SOX-mediated decay is restored on the GFP-SRE* reporter. Since IL-6 is known to also escape decay mediated by closely related SOX-homologs muSOX and BGLF5, we wondered whether the GFP-SRE* would also be susceptible to these endonucleases. As shown in Fig. 2C, a single point mutation at position 74 in the SRE also renders transcripts susceptible to degradation from SOX homologs. Taken together, these data reveal that m^6^A modification of the 3’ UTR of IL-6 promotes its escape from SOX.

### The m6A reader YTHDC2 binds to and promotes the SRE escape from SOX

A previous ChIRP-MS screen had identified a number of host proteins that can bind the SRE element [15]. One of these predicted interactors was the m^6^A reader YTHDC2. Several reports have demonstrated that YTHDC2 directly binds to m^6^A-modified mRNAs often within 3’ UTRs [44-47]. YTHDC2 itself is an RNA helicase and its binding to mRNA has been associated with alteration of RNA stability [44-47]. We first confirmed the interaction between YTHDC2 and SRE-bearing mRNA by performing IPs from cells transfected with a GFP reporter fused to the WT SRE (FIG 3A). In agreement with our previous observations, YTHDC2 binding to the m^6^A deficient SRE* was reduced compared to the WT SRE, confirming that YTHDC2 is recruited to the SRE as an m^6^A reader (FIG 3A).

**Figure 3:**
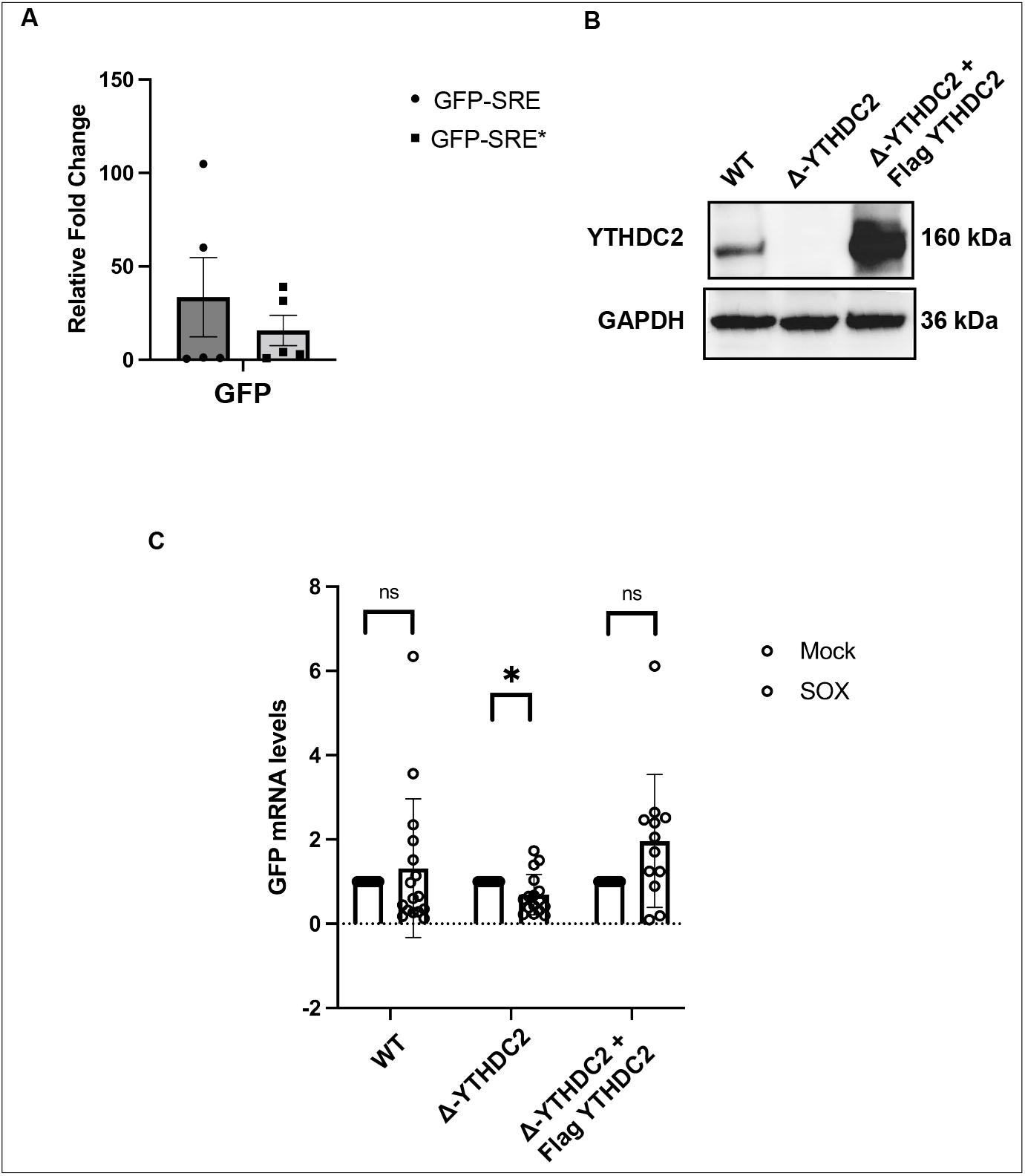
YTHDC2 is necessary for IL-6’s evasion of SOX. **A)** Cells were transfected with Flag-tagged-YTHDC2 and a GFP reporter as indicated. Cells were crosslinked and immunoprecipitated using Flag-coated beads. RNA fraction was collected and used for RT-qPCR. **B)** 293TΔYTHDC2 cells were obtained by stably expressing and single cell selecting 293TCas9 cells expressing a YTHDC2 targeting guide RNA. Cells clones were tested for knock out efficiency by western blot using a YTHDC2 antibody and GAPDH as a loading control. YTHDC2 expression in these cells was rescued by transfecting Flag-tagged-YTHDC2 on a plasmid. **C)** 293TΔYTHDC2 or WT cells were transfected with SOX (or mock) along with a GFP-SRE reporter. RNA was then collected and used for RT-qPCR.

Since the m^6^A modification that we identified on the SRE appears to be important to promote protection from SOX induced degradation, we next asked whether this protective phenotype was being mediated through the recruitment of YTHDC2. We therefore used Cas9-based genome editing to generate YTHDC2 knockout clones in HEK293T cells (now referred to as 293TΔYTHDC2). After confirming knock out efficiency (FIG 3B), we used this cell line to assess how the lack of YTHDC2 expression would affect the SRE stability in the face of SOX-mediated decay. 293TΔYTHDC2 were transfected with our GFP-SRE reporter along with SOX (or mock) and RNA was extracted and used for RT-qPCR (FIG 3C). As expected, SOX does not affect the RNA levels of the GFP-SRE reporter in WT 293T control cells. However, SOX-mediated decay is restored when YTHDC2 expression is knocked down. To ensure that this defect in protection from SOX was not due to off target effects of generating the 293TΔYTHDC2 cell line, we rescued YTHDC2 using ectopic expression (FIG 3B). In these cells, the GFP-SRE stability was rescued to normal levels even in the presence of SOX (FIG 3C). Taken together, these results support a role for the m^6^A reader YTHDC2 in protecting transcripts from SOX degradation.

## Discussion

Herpesviruses extensively manipulate the fate of host transcripts during lytic reactivation through the use of virally encoded endonucleases. In the case of KSHV, the viral endonuclease SOX targets a wide array of mRNAs *via* a sequence specific degron and cleaves around 70% of mRNAs in the cell [1, 48, 49]. This allows the virus unfettered access to the host expression machinery for viral replication. Previous work has shown that among the 30% of transcripts that escape this SOX-mediated decay, there is subset of transcripts that carry an RNA stability element located in their 3’UTR that specifically enables this resistance phenotype against viral but not cellular endoribonucleases known as the SOX-resistant element (SRE) [10, 13-15]. While this escape mechanism remains largely uncharacterized, it is known that this RNA element is not conserved in sequence among escaping transcripts but rather adopts a common RNA structure which has been hypothesized to serve as a protein recruitment platform. Past studies have explored proteins bound to this RNA element and found several m^6^A readers within the SRE RNA-protein complex [15]. We thus hypothesized that the RNA modification m^6^A, which is prevalent and integral to both host and viral transcript fate, may be involved in viral endonuclease escape. This led us to preform m^6^A-eCLIP sequencing on KSHV latent and lytic cells.

The m^6^A-eCLIP confirmed previous results seen where upon reactivation there is an overall decrease in methylation on host transcripts and a massive increase in methylation of viral transcripts [35, 36, 42]. Furthermore, we observed a 5’ UTR hypomethylation and a concomitant 3’ UTR hypermethylation following KSHV reactivation from latency. m^6^A modifications are known to occur mainly co-transcriptionally on the adenines that are located within a DRACH motif by m^6^A writer proteins. It is possible that this shift of m^6^A deposition towards 3’UTR results from alternative spliceforms being expressed during KSHV lytic infection and that possibly, these transcripts have more favorable DRACH motifs. This can be seen in the shift of the types of transcripts being methylated in Fig 1C. Viruses are known to affect the global gene expression landscape and it would thus not be surprising to see that those expressed during lytic infection have alternative 3’UTR favoring m^6^A deposition. This is also in line with our observation that RNA splicing genes are more m^6^A modified during the lytic cycle, which could suggest that they are more solicited and possibly more expressed. However, it is still unknown what dictates the m^6^A writing machinery to prefer one DRACH motif over another. It is possible that upon lytic reactivation a change occurs in the cell that causes m^6^A writing to change its “priority”. This is supported by a couple of genes like GADD45B and ARMC10, whose m^6^A transcript landscape shifts during lytic reactivation as well as the DRACH motifs that are methylated (Fig 1C). Interestingly, we do know that the m^6^A deposition on the IL-6 SRE occurs independently of whether this SRE is in the context of the full transcript. Indeed, our results indicated that the presence of the SRE on a GFP reporter is enough to mediate the same “SOX blocking” effect as in the endogenous mRNA. This suggests that the m^6^A machinery is likely more influenced by a DRACH motif in the proper context and/or presented in the proper structure than other determinants far away from the DRACH motif chosen. Since m^6^A is mainly deposited co-transcriptionally perhaps the difference we see in DRACH motif preference is due to transcriptional rate. Another alternative is given the increase of RNA splicing, m^6^A writers preferentially recognize DRACH motifs in actively spliced RNA as a result.

We were able to show that the m^6^A site in the IL-6 SRE recruits YTHDC2, and further demonstrate that the recruitment of this m^6^A reader is necessary for its protection from SOX. YTHDC2 comes from the YTH family which boast a YTH binding domain to interact with m^6^A directly albeit with low affinity. Interestingly all the other YTH proteins are around 500-750 aa and composed of primarily low complexity disordered regions while YTHDC2 is close to 1,400 aa in length and has several other known domains besides the canonical YTH domain: an R3H, helicase, ankyrin repeats, HA2 and OB-fold domains [44, 45]. Little is known about the function of the canonically cytoplasmic YTHDC2. It has been reported that it may contribute to increased RNA decay by binding select transcripts and XRN1 [45, 46]. Other studies have shown that it enhances translation efficiency unwinding RNA transcripts while bound to the ribosome [44, 50]. This puts it in direct contrast with YTHDC1 which is nuclear and has roles in RNA splicing and chromatin modification [51-53]. YTHDC2 functions more in line with the cytoplasmic YTHDFs 1-3 which have been shown to bind m^6^A containing transcripts and enhance translational activity or mRNA decay [54-57]. What many of the YTH proteins have in common when binding their transcripts is that they function in complex with other proteins. This is consistent with our hypothesis that although YTHDC2 is necessary for the protection of IL-6 from SOX-mediated decay, it is most likely not sufficient. A previous study has shown that IL-6 binds with nucleolin (NCL), HuR and AUF-1 in a protective complex [10, 14, 15]. It is likely that YTHDC2 works in concert with these proteins and possibly others to either occlude SOX targeting *via* their presence or by relocalizing the transcript where SOX cannot target IL-6. There is also a possibility that YTHDC2 helicase function may be necessary for protection and perhaps the unwinding of the IL-6 transcript removes an internal mRNA secondary structure that is essential for SOX targeting. Furthermore, while our data supports the role of m^6^A as an important contributor to SOX resistance, it also emerges that this is not the sole answer of SRE protection. We did not find a consistent pattern in lytic-specific m^6^A deposition in other known or predicted SOX-resistant transcripts. These escaping mRNAs either had no change in their m^6^A status upon lytic infection or had lytic-specific peaks outside of their 3’UTR. This indicates that it is not lytic infection *per se* that triggers this escape phenotype and/or directs m^6^A deposition, but rather that some m^6^A modified mRNAs are compatible with assembling a protective complex against SOX. Therefore, not all m^6^A transcripts turn out to be SOX-resistant which is in line with our observations that only select transcripts among the 20% spared from SOX decay are actively escaping degradation. We thus hypothesize that these m^6^A modifications must be in the proper context and recruit a specific set of protective proteins in order to be active. However, now that we have a clearer idea of what m^6^A reader may be involved in this mechanism, it would be interesting to reverse our question and search for new escapees using either their m^6^A pattern and/or by investigating what transcripts are bound by YTHDC2 during KSHV lytic infection. Furthermore, given that the regulation of RNA fate is a crucial step in hijacking the host cell, it is perhaps unsurprising that several viruses use widespread RNA decay to take over their hosts. It would be interesting to investigate the contribution of m^6^A modifications and YTHDC2 role in these other viral families that also deploy host shutoff as a way to overtake the host.

## Materials and Methods

### Cells and transfections

HEK293T cells (ATCC) were grown in Dulbecco’s modified Eagle’s medium (DMEM; Invitrogen) supplemented with 10% fetal bovine serum (FBS). The KSHV-infected renal carcinoma human cell line iSLK.219 (kind gift from B. Glaunsinger) bearing doxycycline-inducible RTA was grown in DMEM supplemented with 10% FBS [58]. Lytic reactivation cells were induced by the addition of 0.2 μg/ml doxycycline (BD Biosciences) and 110 μg/ml sodium butyrate for 72 h. The 293T□YTHDC2 knockout clone and control Cas9-expressing cells were made by transducing HEK293T cells as previously described [59, 60]. Briefly, lenti-Cas9-blast lentivirus was spinfected onto a monolayer of HEK293T cells, which were then incubated with 20 μg/ml blasticidin for selection of transduced cells. These HEK293T-Cas9 cells were then spinfected with lentivirus made from pLKO-tet on containing the YTHDC2 sgRNA sequence designed using the broad institute analysis tool and checked for off target effects. After selection using and 1 μg/ml puromycin, the pool of YTHDC2 knockout cells was then single-cell cloned in 96-well plates and individual clones were screened by western blot to determine knockout efficiency.

For DNA transfections, cells were plated and transfected after 24h when 70% confluent using PolyJet (SignaGen).

### Plasmids

The GFP-based reporters and SOX expression plasmids were described previously [15]. The SRE* reporter was generated by introducing an A to T point mutation at position 74 of the SRE using the Quickchange site directed mutagenesis protocol (Agilent) using the primers described in Table S1. YTHDC2 expression plasmid was kindly provided by Dr. Chuan He.

### RT-qPCR

Total RNA was harvested using TRIzol according to the manufacturer’s protocol. cDNAs were synthesized from 1 μg of total RNA using AMV reverse transcriptase (Promega) and used directly for quantitative PCR (qPCR) analysis with the SYBR green qPCR kit (Bio-Rad). Signals obtained by qPCR were normalized to those for 18S unless otherwise noted.

### Western blotting

Cell lysates were prepared in lysis buffer (NaCl, 150 mM; Tris, 50 mM; NP-40, 0.5%; dithiothreitol [DTT], 1 mM; and protease inhibitor tablets) and quantified by Bradford assay. Equivalent amounts of each sample were resolved by SDS-PAGE and Western blotted with the following antibodies at 1:1,000 in TBST (Tris-buffered saline, 0.1% Tween 20): rabbit anti-YTHDC2 (Abcam), and mouse anti-GAPDH (Abcam). Primary antibody incubations were followed by horseradish peroxidase (HRP)-conjugated goat anti-mouse or goat anti-rabbit secondary antibodies (1:5,000; Southern Biotechnology).

### MeRIP-qPCR

HEK293T cells were transfected as indicated then total RNA was extracted using TRIzol. Pulldowns were performed using protein G Dynabeads (Invitrogen) with 10ug of m6A antibody (Sigma Aldrich) and 100ug of RNA in MeRIP buffer (50mM Tris-HCl @7.4 pH,150mM NaCl, 1mM EDTA, 0.1% NP-40, millipore H2O) and 1ul of RNAsin (Promega) per sample overnight at 4C. After extensive washing, samples are eluted in MeRIP buffer containing 6.7mM sodium salt for 30mins at 4C. cDNAs were then obtained from 1 μg of total RNA using AMV reverse transcriptase (Promega) and used directly for quantitative PCR (qPCR) analysis with the SYBR green qPCR kit (Bio-Rad).

### RIP

Cells were crosslinked in 1% formaldehyde for 10 minutes, quenched in 125mM glycine and washed in PBS. Cells were then lysed in low-salt lysis buffer [NaCl 150mM, NP-40 0.5%, Tris pH8 50mM, DTT 1mM, MgCl2 3mM containing protease inhibitor cocktail and RNase inhibitor] and sonicated. After removal of cell debris, specific antibodies were added as indicated overnight at 4°C. Magnetic G-coupled beads were added for 1h, washed three times with lysis buffer and twice with high-salt lysis buffer (low-salt lysis buffer except containing 400mM NaCl). Samples were separated into two fractions. Beads containing the fraction used for western blotting were resuspended in 30μL lysis buffer. Beads containing the fraction used for RNA extraction were resuspended in Proteinase K buffer (NaCl 100mM, Tris pH 7.4 10mM, EDTA 1mM, SDS 0.5%) containing 1μL of PK (Proteinase K). Samples were incubated overnight at 65°C to reverse crosslinking. Samples to be analyzed by western blot were then supplemented with 10μL of 4X loading buffer before resolution by SDS-PAGE. RNA samples were resuspended in Trizol and were processed as described above.

### eCLIP and RNA-seq

ISLK.219 cells were harvested in their latent phase or 48h post reactivation. RNA was then extracted by TRIzol and purified as described above. The samples were processed by EclipseBio as described in their user guide, performing 150 Paired End run on NovaSeq6000 on was PolyA selected RNA. Ratio of IP and input reads were evaluated in each cluster, and clusters with IP/input enrichment greater than 8-fold and associated p-value < 0.001 were defined as significant “peaks”. PureCLIP was used to identify m6A sites with a single nucleotide resolution. This algorithm identifies crosslink sites in eCLIP experiments and assess enrichment of DRACH motif relative to reads starts in IP and input libraries, as well as what fraction of identified crosslink sites are positioned on DRACH motifs.

### Statistical analysis

All results are expressed as means ± standard errors of the means (SEMs) of experiments independently repeated at least three times. Unpaired Student’s *t*est was used to evaluate the statistical difference between samples. Significance was evaluated with *P* values as indicated in figure legends.

## Acknowledgments

We thank all members of the Muller lab for helpful discussions and suggestions. We are grateful to Dr. Chuan He for the YTHDC2 expressing plasmid and Britt Glaunsinger for cells and antibodies. We also thank Eclipse Bio for their support.

## Supplemental Figures

**Figure S1:** qPCR of KSHV transcripts.

**Figure S2:** qPCR of MeRIP control transcripts, C19ORF66 as a negative control and DiCER as the positive control

**Table S1:** Primer dataset. Contains sequencing and qPCR primers used.

**Table S2:** M6A-eCLIP IP iSLK-Lat Crosslink Site dataset. Crosslink sites were identified using the tool <u>PureCLIP</u>. Input samples were used as a background control for crosslink site identification. The score is the log posterior probability ratio of the first and second likely state. The fold change is the fold change of IP read starts vs. input read starts.

**Table S3:** M6A-eCLIP IP iSLK-Lyt Crosslink Site dataset. Crosslink sites were identified using the tool <u>PureCLIP</u>. Input samples were used as a background control for crosslink site identification. The score is the log posterior probability ratio of the first and second likely state. The fold change is the fold change of IP read starts vs. input read starts.

